## Supplementary material for "Master Regulator Analysis of the SARS-CoV-2/Human interactome": Table S2

| human&pangolin | bat | human&pangolin | bat | human&pangolin | bat |
|---|---|---|---|---|---|
| ALA16 | THR16 | HIS228 | ARG228 | LEU568 | LYS568 |
| GLU23 | ASP23 | SER254 | PHE254 | ALA569 | THR569 |
| THR27 | MET27 | ALA296 | GLU296 | VAL573 | ILE573 |
| GLU35 | LYS35 | VAL298 | LEU298 | GLY575 | ASP575 |
| TYR41 | HIS41 | GLN325 | GLU325 | LYS577 | ARG577 |
| ASN49 | ASP49 | GLU329 | ASN329 | ARG582 | GLY582 |
| THR55 | ASN55 | ASP367 | GLU367 | ASN586 | LYS586 |
| ASN63 | ASP63 | ILE407 | VAL407 | LYS596 | GLN596 |
| GLN86 | GLU86 | ALA412 | VAL412 | LYS600 | ARG600 |
| ALA99 | ILE99 | ILE421 | MET421 | ASN601 | LYS601 |
| SER106 | PRO106 | THR445 | ASN445 | PHE603 | TYR603 |
| THR118 | SER118 | GLN472 | GLU472 | ALA614 | SER614 |
| THR122 | ALA122 | GLU483 | LYS483 | LYS631 | ASN631 |
| ASN134 | LYS134 | TYR521 | PHE521 | VAL658 | GLU658 |
| ASN137 | LYS137 | GLN522 | GLU522 | ARG671 | TRP671 |
| SER170 | ALA170 | GLN526 | HIS526 | PHE684 | HIS684 |
| ALA193 | GLY193 | GLN531 | ARG531 | LYS689 | GLY689 |
| ASN194 | TYR194 | LYS534 | GLN534 | VAL691 | LEU691 |
| GLY205 | ARG205 | GLU536 | ASP536 | GLY751 | ALA751 |
| GLY211 | GLU211 | GLU549 | ASP549 | PHE762 | ILE762 |
| TYR215 | PRO215 | GLN552 | LYS552 | LYS769 | ARG769 |
| ILE223 | MET223 | LEU560 | VAL560 | LYS771 | THR771 |
| GLU224 | LYS224 | PRO565 | ALA565 | ALA782 | SER782 |
