## Supplementary figures and images for "Master Regulator Analysis of the SARS-CoV-2/Human interactome"

### Figure S1

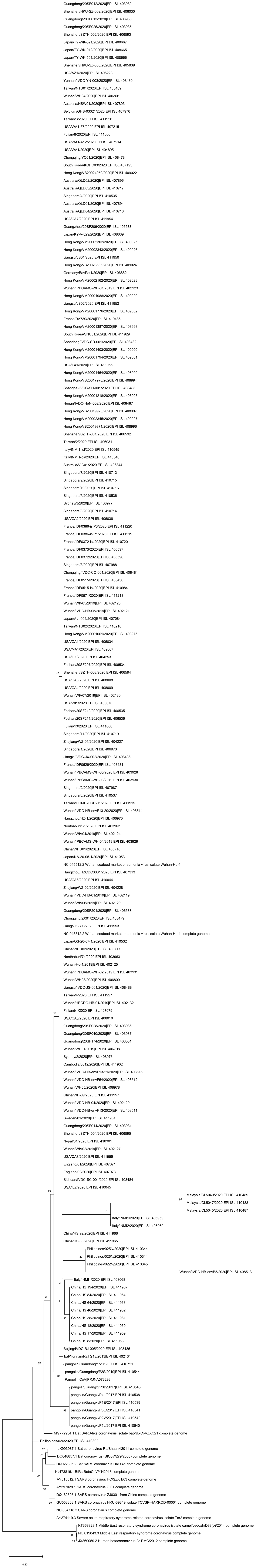
